# ROGUE: an entropy-based universal metric for assessing the purity of single cell population

**DOI:** 10.1101/819581

**Authors:** Baolin Liu, Chenwei Li, Ziyi Li, Xianwen Ren, Zemin Zhang

**Author notes:** These authors contributed equally: Baolin Liu and Chenwei Li.

## Abstract

Single-cell RNA sequencing (scRNA-seq) is a versatile tool for discovering and annotating cell types and states, but the determination and annotation of cell subtypes is often subjective and arbitrary. Often, it is not even clear whether a given cluster is uniform. Here we present an entropy-based statistic, ROGUE, to accurately quantify the purity of identified cell clusters. We demonstrated that our ROGUE metric is generalizable across datasets, and enables accurate, sensitive and robust assessment of cluster purity on a wide range of simulated and real datasets. Applying this metric to fibroblast and B cell datasets, we identified additional subtypes and demonstrated the application of ROGUE-guided analyses to detect true signals in specific subpopulations. ROGUE can be applied to all tested scRNA-seq datasets, and has important implications for evaluating the quality of putative clusters, discovering pure cell subtypes and constructing comprehensive, detailed and standardized single cell atlas.

## Introduction

Tissues are complex milieus comprising various cell types and states with specialized roles^1^. Characterizing the property and function of each pure cell type is a long-standing challenge in biological and medical disciplines. The recent advances in scRNA-seq have transformative potential to discover and annotate cell types, providing insights into organ composition^2^, tumor microenvironment^3^, cell lineage^4^ and fundamental cell properties^5^. However, the identification of cell clusters is often determined by manually checking specific signature genes, which are arbitrary and inherently imprecise. In addition, different methods and even parameters used for normalization, feature selection, batch correction and clustering can also confound the final identified clusters^6^, thus motivating the need to accurately assess the purity or quality of identified clusters (Fig. 1a).

**Fig. 1.**
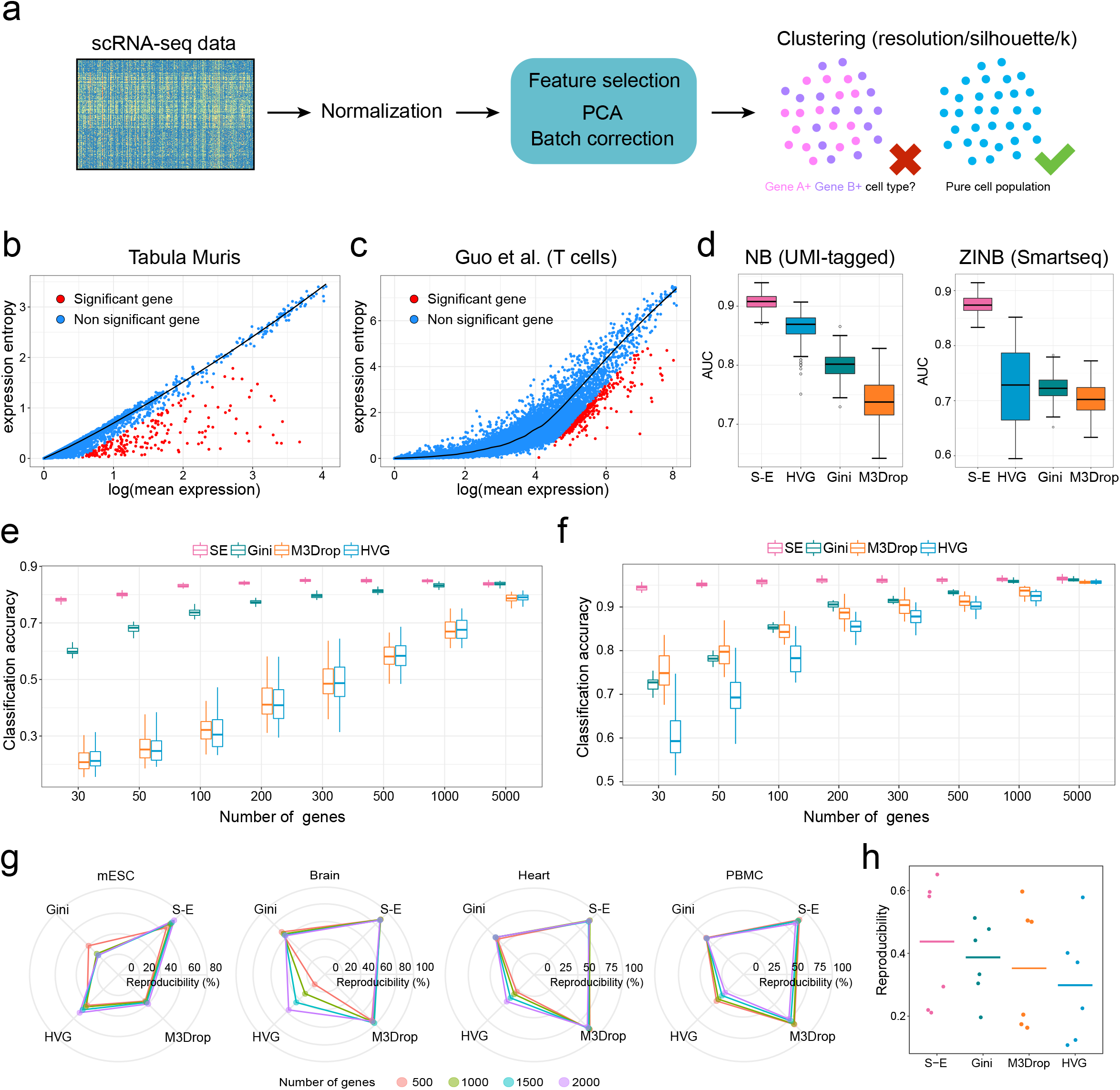
The expression entropy model. **(a)** Identifying pure cell subtypes in unsupervised single-cell data analysis. **(b)** The *S* - *E* plot of the Tabula Muris (droplet) dataset. Each point represents one gene. The relationship between *S* and *E* was fitted with LOESS regression for each gene. **(c)** The *S*-*E* plot of a T cell dataset^21^ obtained by Smart-seq2 protocol. **(d)** Accuracy in identifying differentially expressed genes on data simulated from both NB (left) and ZINB (right) distribution, with subpopulation containing 50% of the cells. The center line indicates the median AUC value of n=50 repeated runs. The lower and upper hinges represent the 25th and 75th percentiles respectively, and whiskers denote 1.5 times the interquartile range. **(e,f)** Discriminating power of genes selected by *S*-*E* model, Gini, M3Drop and HVG estimated by RF with 50 times cross-validation on both droplet-based dataset **(e)** and full-length based dataset **(f)** listed in Supplementary Table 1. The classification accuracy was measured as the percentage of query cells that were assigned the correct label. **(g,h)** Reproducibility of features across four groups of replicates **(g)** and four human pancreas datasets **(h)** listed in Supplementary Table 3. Each point in **(h)** denotes a pair of datasets, and horizontal lines represent the median values across all pairs.

A pure cluster here is defined as a population where all cells have identical function and state without variable genes. The importance of purity assessment is particularly relevant for analyses that aim to discover novel pure subtypes and further detect the true biological signals. For example, signature genes specific to a pure subpopulation may be mistakenly considered as the common signals of a mixture due to less guided clustering and annotation. The purity evaluation could therefore eliminate such misleading conclusions, potentially aiding our understanding of cellular function, state and behavior. While pioneering approaches such as silhouette^7^, DendroSplit^8^, and distance ratio^9^ have been devoted to determining the optimal number of identified clusters by calculating the ratio of within-cluster to inter-cluster dissimilarity, they are not comparable among datasets and have poor interpretability of cluster purity. For example, an average silhouette value of 0.7 indicates a fairly strong consistency for a given cluster, but it is still unknown whether this cluster is a pure population or a mixture of similar subpopulations especially when frequent dropout events occur.

The challenges presented by purity evaluation can be broadly addressed by investigating the number of “infiltrating” non-self cells and variable genes, which are suited to the intended areas of unsupervised variable gene detection. Given its importance, diverse methods^10^ have been proposed for the quantification and selection of highly variable genes. In particular, scran^11^ aims to identify variable genes by comparing variance to a local regression trend. However, the over-dispersion, coupled with the high frequency of dropout events, would often result in “swamping” of useful information, causing the deterioration of the results of such variance-based approaches^12^. Alternatively, Gini coefficient^13^ could be used to quantify the variation in gene expression, but the limited performance restricts its scalability. New probabilistic approaches for variable gene selection using dropout rates have also been recently adapted^14^, with the advantage of supporting both pseudotime analysis and discrete clustering, but their usage of dropout metric hinder the capturing of the global distribution of gene expression. Although highly informative genes can also be determined by inspecting their weights during multiple iterations of dimensionality reduction^15^, such ad hoc approaches are computationally intensive and do not provide independent metrics for gene expression variability.

Here, we present an entropy-based model to measure the randomness of gene expression in single cells, and demonstrate that this model is scalable across different datasets, capable of identifying variable genes with high sensitivity and precision. Based on this model, we propose the ROGUE statistic to quantify the purity or homogeneity of a given single cell population while accounting for other technical factors. We demonstrate that the ROGUE metric enables accurate and unbiased assessment of cluster purity, and thus provides a universal measure to evaluate the quality of both published and newly generated cell clusters. Applying ROGUE to B cell and fibroblast analyses, we identified additional pure subtypes and demonstrate the application of ROGUE-guided analysis in detecting the real biological signals. Our approach is broadly applicable for any scRNA-seq data, and is implemented in an open-source R package ROGUE (https://github.com/PaulingLiu/ROGUE), which is freely available.

## Results

### Expression entropy model enables sensitive and accurate identification of variable genes

As scRNA-seq data can be approximated by negative binomial (NB) or zero-inflated NB (ZINB) distribution^16,17^, we considered the use of the statistic, *S* (*expression entropy* — differential entropy of expression distribution, as defined in Methods), to capture the degree of disorder or randomness of gene expression. Notably, we observed a strong relationship between *S* and the mean expression level (*E*) of genes, thus forming the basis for our expression entropy model (*S*-*E* model, Fig. 1b-c). Moreover, *S* is linearly related to *E* for the Tabula Muris dataset^2^ as expected (Fig. 1b), which is characteristic of current droplet experiments, hence demonstrating the NB nature of UMI-based datasets (Methods). For a heterogeneous cell population, certain genes would exhibit expression deviation in fractions of cells, leading to constrained randomness of its expression distribution and hence the reduction of *S*. Accordingly, informative genes can be obtained in an unsupervised fashion by selecting genes with maximal *S*-reduction (*ds*) against the null expectation (Methods).

To illustrate the performance of *S*-*E* model, we benchmarked our method against HVG^11^, Gini^13^ and M3Drop^14^ on data simulated from both NB and zero-inflated NB (ZINB) distribution (Methods). For a fair comparison, we generated a total of 1600 evaluation datasets and used AUC as a standard to test the performance of each method. As a tool to identify genes specific to rare cell types, Gini outperformed HVG and M3Drop when there were subpopulations accounting for less than 20% of the cells. In contrast, HVG was slightly better than Gini and M3Drop in the presence of cell subpopulations with a larger proportion. Notably, expression entropy model consistently achieved the highest AUC score and significantly outperformed other gene selection methods in all tested cases with varied subpopulation proportions or gene abundance levels (Fig. 1d and Supplementary Figs. 1 and 2).

**Fig. 2.**
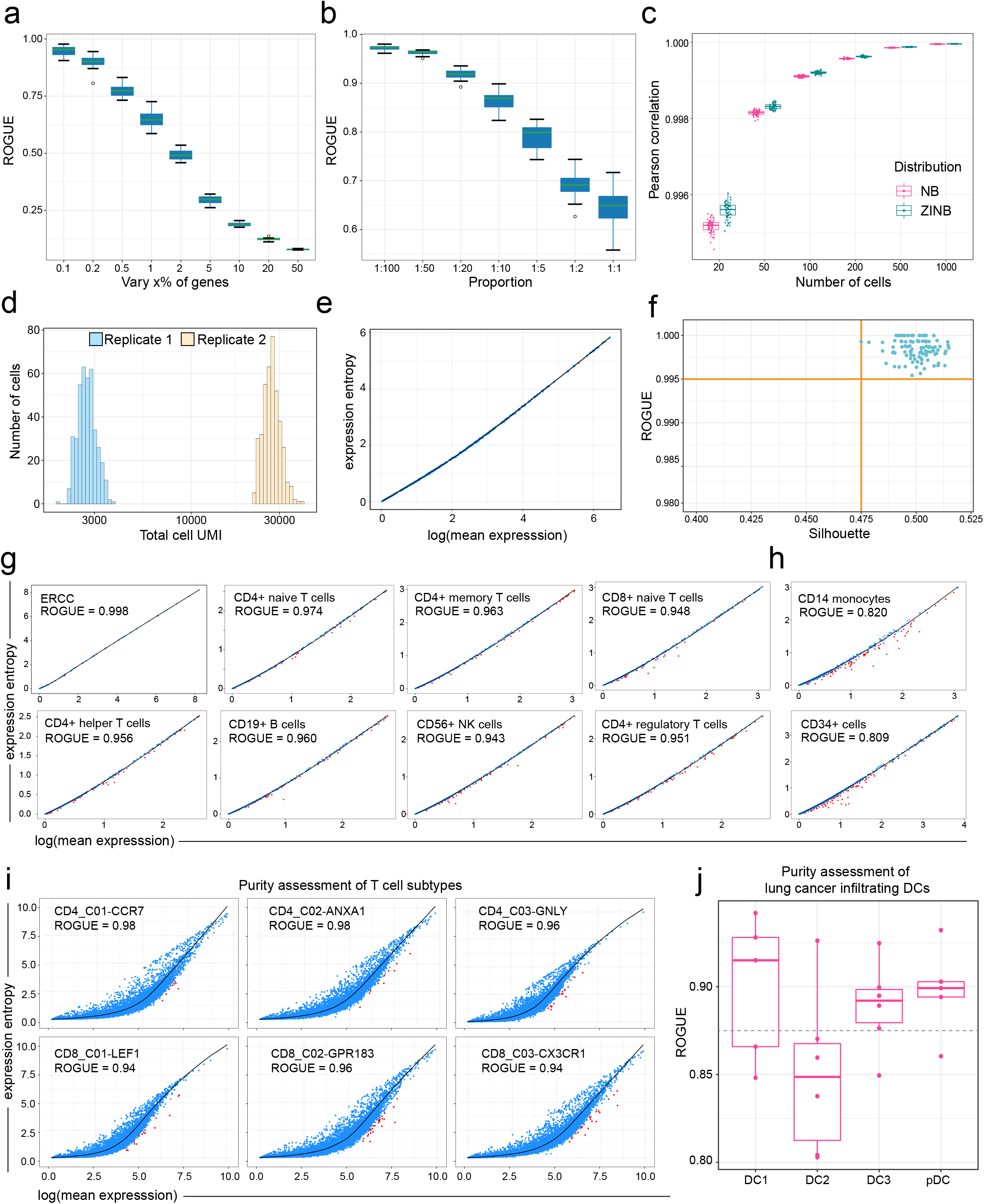
ROGUE use and performance. **(a)** The ROGUE index decreases monotonically with increasing varied genes in each simulated mixture consisting of two cell types (1:1). The center line indicates the median value of n=50 repeated ROGUE runs. The lower and upper hinges represent the 25th and 75th percentiles respectively, and whiskers denote 1.5 times the interquartile range. **(b)** The ROGUE values for the simulated mixtures with cell type sizes ranging from 1:100 to 1:1. In each mixture, the number of varied genes was 1% of the total gene number (n = 20,000). **(c)** Pearson correlations of *S* between the randomly down-sampled datasets (n=50 runs for each) and the entire datasets (2,000 cells) simulated from both NB and ZINB distribution. **(d)** Sequencing depth distribution (total UMI counts/cell) for two simulated replicates. The replicate 2 has a sequencing depth 10 times that of replicate 1. **(e)** The *S*-*E* plot of the mixture of replicates 1 and 2 shown in **(d)**. **(f)** ROGUE values of n=100 mixtures versus the silhouette values for every two replicates within individual mixtures. A high silhouette value indicates a substantial difference in sequencing depth between two replicates. **(g**,**h)** The *S*-*E* plots and corresponding ROGUE values of 10 cell populations from the PBMC dataset^20^. **(i)** Purity assessment of six human T cell populations. **(j)** Purity evaluation of lung cancer infiltrating DCs, with each point representing a patient.

To validate our unsupervised feature selection method in real datasets, we performed cross-validation experiments using random forest classifier (RF)^18^, with 70% cells from the original sample as reference and remaining 30% cells as query set (Methods). Intuitively, gene sets that enable higher classification accuracy are more biological meaningful. Using 14 previously published datasets derived from both droplet-based and full-length protocols (Supplementary Table 1), we demonstrated that our method consistently identified genes with greater ability of classification when different number (30-5000) of genes were selected, followed by Gini, HVG and M3Drop (Fig. 1e,f and Supplementary Figs. 3 and 4). Specially, our *S*-*E* model showed notable superiority when fewer genes (30-100) were used, demonstrating its sensitivity. Taken together, these results suggest that genes identified by our model are more informative and biologically discriminating.

**Fig. 3.**
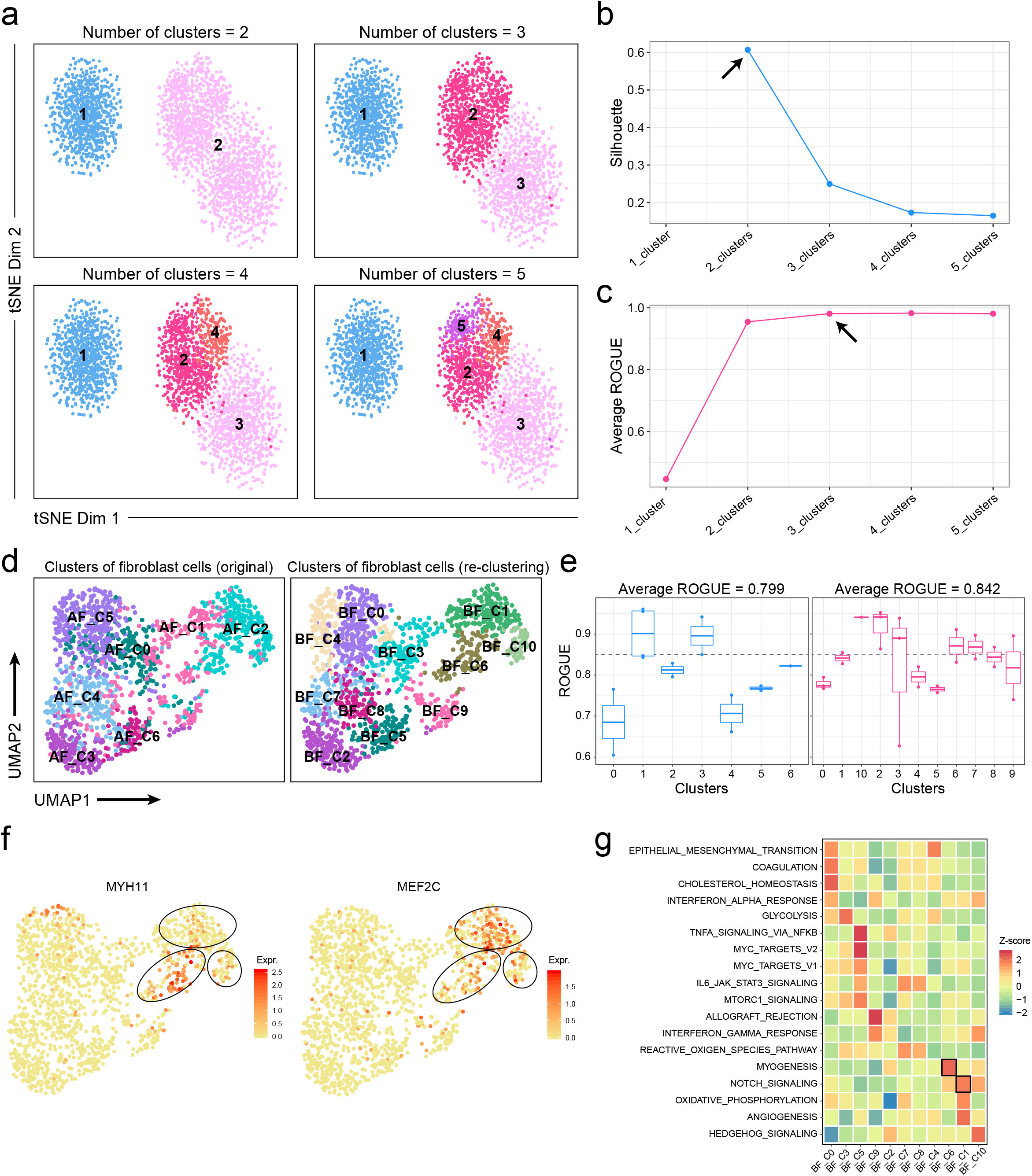
ROGUE enhances single cell clustering and cell type identification. **(a)** t-SNE plots of a simulated dataset containing three cell types. **(b**,**c)** Corresponding silhouette values **(b)** and average ROGUE values **(c)** when there were 2, 3, 4 and 5 putative clusters respectively. **(d)** UMAP plots of lung cancer associated fibroblasts, color-coded by clusters in original paper (left; Supplementary Fig. 9a) and re-clustered labels (right). **(e)** ROGUE values of different clusters before (left) and after (right) re-clustering. Each point represents a patient. **(f)** UMAP plot of expression levels of MYH11 and MEF2C. **(g)** Differences in hallmark pathway activities scored using GSVA.

**Fig. 4.**
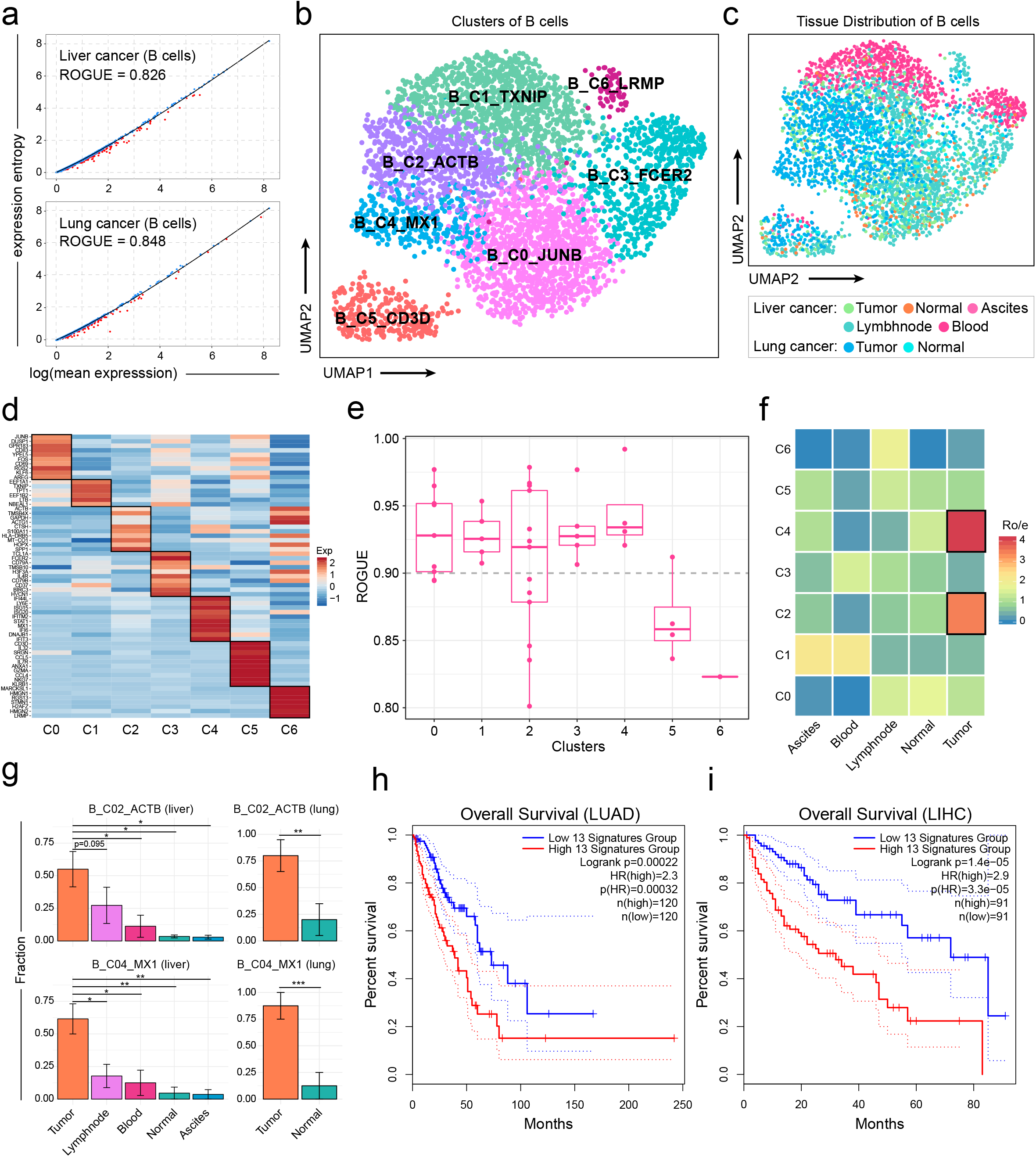
ROGUE-guided analysis in the identification of pure B cell subtypes. **(a)** The *S*-*E* plots and ROGUE values of liver and lung tumor-infiltrating B cells, respectively. **(b,c)** UMAP plots of 4,291 B cells, color-coded by their associated clusters **(b)** and tissues **(c)**. **(d)** ROGUE values of seven identified B cell subtypes. Each point represents a patient. **(e)** Gene expression heatmap of 7 B cell clusters. Rows denote marker genes and columns denote different clusters. **(f)** Tissue preference of each B cell subtype in liver cancer estimated by R_O/E_^21^, the ratio of observed to expected cell numbers calculated by chi-square test. **(g)** The fractions of B_C02_ACTB and B_C04_MX1 in each patient across tissues. *p<0.05, **p<0.005, Student’s t test. **(h,i)** The Kaplan–Meier curves of TCGA LUAD **(h)** and LIHC **(i)** patients grouped by the 13 markers (Supplementary Table 5) of B_C02_ACTB.

Since datasets derived from the same biological system are expected to have reproducible informative genes^14^, we tested how our expression entropy model behave using technical replicates from different tissues (Supplementary Table 2). Notably, genes identified by our *S*-*E* model were more reproducible when top 500-2000 genes were used (Fig. 1g). In addition, we also considered four pancreatic datasets (Supplementary Table 3) derived from different technologies and labs. These real datasets are more complex than technical replicates as they included systemic nuisance factors such as batch effects. Despite substantial systematic differences, our model consistently achieved the highest reproducibility score (Fig. 1h), demonstrating that *S* - *E* model provides reliable, robust and accurate identification of informative genes, hence forming the foundation for the subsequent purity assessment of single cell clusters.

### The ROGUE index for quantifying the purity of cell population

Unsupervised clustering is currently the standard method to identify new cell types^19^. However, the identification and annotation of a putative subtype is often arbitrary and lack a quantitative metric for robust purity evaluation. Here, we take advantage of the wide applicability of *S*-*E* model to scRNA-seq data and introduce the statistic ROGUE (Ratio of Global Unshifted Entropy) to measure the purity of single cell populations (Methods). Intuitively, a cell population with no significant *ds* for all genes will receive a ROGUE value of 1, indicating it is a completely pure subtype or state. In contrast, a population with maximum summarization of significant ds will yield a purity score of ~0.

We investigated the performance of ROGUE on 1860 cell populations simulated from both NB and ZINB distribution (2000 cells × 20000 genes each), with 0.1%-50% genes varied in a second cell type (Methods). A cell population with both fewer “infiltrating” non-self cells and varied genes would yield a high purity score, while a population with converse situation is expected to yield a low purity score. It is evident that the ROGUE index decreased monotonically with the heterogeneity of cell populations (Fig. 2a,b and Supplementary Figs. 5 and 6). ROGUE performed well even when cell populations contained few varied genes (<1%) and “infiltrating” cells (<1%), indicating ROGUE index provides a sensitive and unbiased measure in response to the degree of cell population purity. To address the potential concern that the number of cells may represent an intrinsic challenge for *S* and ROGUE calculation, particularly if only few cells are collected from given samples, we performed down sampling analysis to test how *S* was impacted by cell numbers. By calculating the Pearson correlations of *S* between the randomly down-sampled datasets and the entire datasets, we found the similarity values of >0.99 and demonstrated that our *S* and ROGUE calculation would not be unaffected by variation in cell number (Fig. 2c).

Sequencing depth can vary significantly across cells, with variation potentially spanning orders of magnitude^2^, and hence contributes to a substantial technical confounder in scRNA-seq data. We sought to investigate whether ROUGE index can accurately assess the purity of single cell population while accounting for this technical effect. As test cases, we simulated increasing molecular counts (sequencing depth) in a second “mock” replicate, with the fold change of gene expression means ranging from 2 to 100 (Fig. 2d and Methods). Despite the substantial technical effect, the mixture of each two simulated replicates is expected to be a pure cell population. Here we used silhouette to measure the degree of replicate-to-replicate differences. The results revealed ROGUE values of ~0.99-1 for each population consisting of two replicates, with silhouette values ranging from 0.25 to 0.75 (Fig. 2e,f and Supplementary Fig. 7a). Thus, ROGUE not only offers a robust and sensitive way to estimate the purity of single cell population, but also accounts for the variation in sequencing depth.

### The ROGUE accurately assesses the purity of cell populations

To illustrate the applicability of ROGUE index to real data, we first considered an ERCC (External RNA Controls Consortium) dataset^20^ with no biological variability, which is an ideal case of pure cell population. This ERCC dataset achieved ROGUE ~1 with only one significant gene. Further, we investigated the fresh peripheral blood mononuclear cells (PBMCs) enriched from a single healthy donor^20^. The authors provided multiple cell types purified by fluorescence-activated cell sorting (FACS), and thus representing a suitable resource for purity assessment. These cell types in Fig. 2g, including CD4/CD8 naïve T cells and CD4 memory T cells, have been shown to be highly homogeneous populations^21^, and were detected high ROGUE values (0.94-1) as expected. In contrast, both CD14 monocytes and CD34+ cells are mixtures of diverse subtypes^20^ and received relatively low ROGUE values (~0.8; Fig. 2h), thus confirming their heterogeneity.

In addition to highly controlled datasets, it is also instructive to investigate how ROGUE index performs on pure subtypes identified by unsupervised clustering. Here we first considered six well defined T cell subtypes from human colorectal cancer^5^, which were generated via the Smart-seq2 protocol. All these pure subtypes achieved high ROGUE values of >0.9 (Fig. 2i), versus 0.78 for complete data (Supplementary Fig. 7b). We next examined four dendritic cell (DC) subsets collected from human lung cancers^22^ and sequenced with inDrop platform. Specially, tumor-infiltrating DC2 cells have been proven to be highly heterogeneous populations^23,24^ and deviated substantially from the other homogeneous cell types including DC1, LAMP3+ DC and pDC (Fig. 2j). Taken together, these results illustrate that our ROGUE represents an effective and direct quantification of cell population purity without being affected by technical characteristics.

### ROGUE-guided analysis enhances single cell clustering and cell type identification

We next evaluated the potential for ROGUE to guide clustering analysis with silhouette, which investigates whether a certain clustering has maximized inter-cluster dissimilarity and minimized within-cluster dissimilarity. As a test case, we simulated a scRNA-seq dataset consisting of three cell types A, B and C (see Methods for details), where cell type A and B were similar subtypes with 1% varied genes. We clustered this dataset into 2, 3, 4, and 5 subpopulations respectively by adjusting the resolution parameter in Seurat^25^ (Fig. 3a), then evaluated the results by inspecting corresponding silhouette and average ROGUE values. Proper clustering of this dataset should result in three subpopulations, one for each cell type. However, silhouette received the maximum value when cell type A co-clustered with B (Fig. 3b), *i.e.*, when only two clusters were identified, suggesting that such measure is poorly interpretable for cluster purity as opposed to ROGUE, which reached saturation when there were three clusters (Fig. 3c). Repeating the simulation with varied differences in cell type A, B and C yielded equivalent performance for these two methods (Supplementary Fig. 8a-f). Since ROGUE can provide direct purity quantification of a single cluster and is independent of methods used for normalization, dimensionality reduction and clustering, it could also be applied to guide the splitting (re-clustering) or merging of specific clusters in unsupervised clustering analyses.

To test how ROGUE could help the clustering of real datasets, we examined a previously reported dataset of cancer-associated fibroblasts (CAFs)^26^, which were collected from lung tumors. CAFs have been reported to represent a highly heterogeneous population and may play a tumor-supportive role in the tumor microenvironment^27^. We found that the 7 identified fibroblast clusters received low ROGUE values (Fig. 3d,e and Supplementary Fig. 9a). We therefore performed re-clustering analysis with the goal of exploring the extent of heterogeneity and identified a total of 11 clusters with a higher average ROGUE value (Fig. 3d,e). In addition to the two classical subtypes of CAFs (myofibroblastic CAFs and inflammatory CAFs), we also found the presence of antigen-presenting CAFs (apCAFs) that was characterized by the high expression of CD74 and MHC class II genes (Supplementary Fig. 9b). The apCAFs were firstly discovered as a fibroblast subtype in mouse pancreatic ductal adenocarcinoma (PDAC), but barely detectable in human PDAC without forming a separate cluster^28^. The considerable existence of apCAFs in lung cancer thus may indicate potential differences between different cancer types.

Furthermore, we noted that the myCAFs (AF_C02_COL4A1, ROGUE=0.81) identified by original authors could be further segregated into three distinct subpopulations, including BF_C01_RGS5 (ROGUE=0.87), BF_C02_ACTA2 (ROGUE=0.84) and BF_C03_GPX3 (ROGUE=0.94). Interestingly, the signature genes of AF_C02_COL4A1 described by original authors were actually specific to one of these three subpopulations, including MEF2C in BF_C01_RGS5 and MYH11 in BF_C02_ACTA2 (Fig. 3f). Pathway analysis also revealed that the NOTCH signaling was activated in BF_C01_RGS5 (Fig. 3g) rather than a common signal of AF_C02_COL4A1^26^. Despite the considerable increase of overall ROGUE index, BF_C00_AOL10A1, BF_C04_COL1A2 and BF_C05_PLA2G2A still received relatively low ROGUE values, thus deserving further investigation. Overall, ROGUE-guided analysis not only discovered novel cell subtypes, but also enabled the detection of the true signals in specific pure subpopulations.

### ROGUE-guided analysis identified pure B cell subtypes in liver and lung cancer

B cells are key components in tumor microenvironment but have unclear functions in antitumor humoral response^29^. Here we investigated previously reported liver and lung tumor infiltrating B cells^26,30^ and found that they received relatively low ROGUE values (Fig. 4a). Thus, we applied further clustering analysis coupled with ROGUE to these B cells in an attempt to discover pure subtypes. A total of 7 clusters were identified, each with its specific marker genes (Fig. 4b-d). Cells from the first B cell subset, B_C0_JUNB, specifically expressed signature genes including JUNB and FOS, thus representing activated B cells^31^. The second subset, B_C1_TXNIP, showed high expression of glycolysis pathway genes (Supplementary Table 4), indicating its metabolic differences. ACTB, a gene involved in antigen presenting, was highly expressed in the third subset (B_C2_ACTB). Pathway activity analysis also revealed a strong antigen processing and presentation signal in this subset (Supplementary Table 4). The fourth cluster, B_C3_FCER2, characterized by high expression of HVCN1 and genes involved in B cell receptor signaling pathway (Supplementary Table 4), was largely composed of pre-activated B cells^30^. The fifth cluster, B_C4_MX1, predominantly composed of interferon induced B cells^32^, expressed high levels of MX1, IFI6 and IFI44L. The sixth cluster, B_C5_CD3D, expressed key markers of both T and B cell lineages (Fig. 4d), thus maybe the dual expressers (DEs)-like lymphocytes^33^ or doublets. The remaining B cells, falling into the seventh cluster, B_C6_LRMP, exhibited high expression of LRMP and RGS13, indicative of the identity of germinal center B cells^34^.

Both DEs/doublets-like and germinal center B cells exhibited low ROGUE values (Fig. 4e), but the limited cells did not permit further clustering. For germinal center B cells, we readily detected the high expression of proliferating marker genes, including MKI67 and STMN1 (Supplementary Fig. 10), in a fraction of these cells, thus explaining the heterogeneity to some extent. In contrast to these two clusters, we found ROGUE values of >0.92 for each of the remaining five clusters (Fig. 4e), demonstrating that they were all highly homogeneous B cell subtypes. By calculating the ratio of observed to expected cell numbers with chi-square test (R_O/E_), we noted that both B_C02_ACTB and B_C04_MX1 contained mainly cells from tumor, with R_O/E_ values > 1 (Fig. 4f). Similar analyses stratified by patient further confirmed this trend (Fig. 4g). Based on the independent TCGA lung adenocarcinoma (LUAD) cohort dataset, patients with higher expression of the marker genes of B_C02_ACTB (normalized by MS4A1; Supplementary Table 5) showed significantly worse overall survival (Fig. 4h). Such survival difference was also observed in TCGA liver hepatocellular carcinoma (LIHC) cohort dataset (Fig. 4i). Thus, the clinical implication deserves further study to investigate what specific roles B_C02_ACTB cells play in tumor microenvironment. In summary, identifying pure subtypes with ROGUE-guided analysis could enable a deeper biological understanding of cell state and behavior.

## Discussion

Purity assessment of identified cell clusters is paramount to the interpretation of scRNA-seq data. This assessment is especially pertinent as increasingly rare and subtle cell subtypes are being uncovered. To address this computational challenge, we present the *S*-*E* model and demonstrate that this model is capable of identifying variable genes with high sensitivity and precision, and thus could be applied to both clustering and potentially pseudotime analyses. By taking advantage of the wide applicability of *S*-*E* model, we develop the statistic ROGUE to quantify the purity of single cell populations. Through a wide range of tests, we demonstrate that our entropy-based measure, ROGUE, is generalizable across datasets from different platforms, protocols and operators, and able to successfully quantify the purity of single cell populations regardless of uncontrollable cell-to-cell variation.

When using ROGUE to assess the purity of four DC subtypes from human lung tumors, we found that DC2 was a heterogeneous population, which is consistent with previous findings^24^. Such heterogeneous populations like DC2 may have different properties and specialized roles in the cancer microenvironment, and could be assessed in a similar fashion with ROGUE. Accordingly, future studies could focus on these cell populations and hence may deepen our understanding of cellular origins of cancer. In addition, ROGUE addresses an important need in unsupervised single cell data analyses, *i.e.*, to effectively assess the quality of published or newly generated clusters. Often, unsupervised clustering may lead to under- or over-clustering of cells due to the lack of universal stands for clustering quality. By quantifying cluster purity with ROGUE before and after clustering or re-clustering, we were able to detect low-purity clusters and perform further analysis to discover pure subtypes. Improving the purity and credibility of the ever-increasing number of cell types is a mounting challenge with explosive efforts towards single cell sequencing, and ROGUE could become a potential universal standard for judging the quality of cell clusters.

Our ROGUE-guided analysis on fibroblasts identified a novel subpopulation in lung cancer, apCAFs, which highly expressed CD74 as well as MHC class II genes and had a strong antigen-presenting signal. These cells have been speculated to deactivate CD4 T cells and decrease the CD8+ to Treg ratio in mouse PDAC^28^, but have unclear role in the lung cancer microenvironment, hence requiring further investigation. Moreover, when applying ROUGE to B cell analysis, we found an interesting pure cluster B_C02_ACTB that displayed high expression of genes involved in antigen processing and presentation. Cells from this cluster were preferentially enriched in tumors and were associated with poor prognostic outcomes in both lung and liver cancer. We therefore hypothesize that these cells may contribute to immune suppression in the cancer microenvironment and hence curtail anti-tumor immunity, although further studies are required to define the roles of these cells. Such approaches for discovering novel or additional pure subtypes can also be extended to other published or newly generated scRNA-seq datasets.

When determining the purity of cell clusters, we recommend a ROGUE value of 0.9 as a suitable threshold, at which the number of “infiltrating” cells and varied genes is well constrained. But for low-quality data or continuous data, the threshold could be determined by considering the global ROGUE values. Although ROGUE can be very efficient and effective, we anticipate that additional extensions could enable enhanced performance, for example, assessing the purity of integrated cell populations from different protocols and platforms. Overall, our ROGUE metric provides a robust, direct and universal measure for cluster purity in the presence of substantial technical confounders. We expect the ROGUE metric to be broadly applicable to any scRNA-seq datasets, and anticipate that our strategy will improve the rigor and quality of unsupervised single-cell data analysis.

## Methods

### Expression entropy model

For droplet datasets, the observed UMI count can be modeled as a NB random variable, which also arises as a Poisson-gamma mixture^35^:

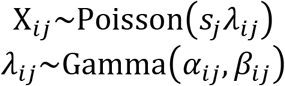

where *λ*_*ij*_ represents the true exp ression value that underlies the observed UMI count X_*ij*_ of gene *i* in cel l *j*, and *S*_*j*_ denotes the size normalization factor in cell *j*. The *α*_*ij*_ and *β*_*ij*_ are shape parameter and rate parameter respectively. Given the assumption that the shape parameter *α* is a constant across cells and genes, and that the rate parameter *β* is a constant of gene *i* across cells^35,36^, *α*_*ij*_ and *β*_*ij*_ can be expressed as *α* and *β*_*i*_, respectively. Then the distributions can be recognized as: *λ*_*i*_~Gamma(*α*, *β*_*i*_) and X_*ij*_~Poisson(*s*_*j*_*λ*_*i*_). We denote 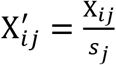 as normalized expression and use 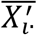 (the mean 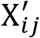 across cells) as the moment estimation of *λ*_*i*_. For the Gamma distribution, the rate parameter could therefore be calculated based on the maximum likelihood estimation:

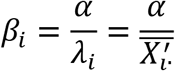

Here we sought to use differential entropy to capture the degree of disorder or randomness of gene expression as we adapted in our supervised gene selection method E-test^37^. For the gamma distributed random variable *λ*_*i*_, its differential entropy can be computed as:

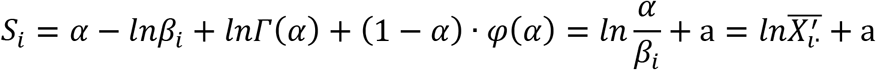

where *φ* is the digamma function and a = α − ln*α* + lnΓ(α) + (1 − α) · φ(α) is a constant. Although other pioneering methods such as Scnorm^38^, scran^39^ and BASiCS^40^ can be used to calculate size factors, we considered the library size normalization defined as^35^:

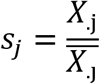

where *X*_.j_ is the total UMI counts in cell *j* and 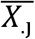 is the mean library size across cells. Accordingly, we can derive that the mean of *s* across cells is 1. Given that the library size is a random variable independent of gene and cell^36^, we can derive that:

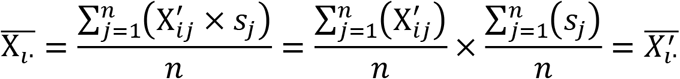

where 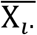 is the observed mean expression of gene *i*, in correspondence to *E*_*i*_ described in the main text. For each cell type, the differential entropy of *λ*_*i*_ could be computed as:

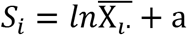

We formulate the null hypothesis that there is only one Poisson-gamma component for each gene in a given population (*H*_0_) and thus the corresponding differential entropy can be calculated with 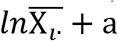. Then we assume that each cell represents its own “cluster” and use *X*_*ij*_ as a moment estimation of the mean expression of such “cluster”. In this way, we define the entropy reduction of gene *i* across *n* cells as:

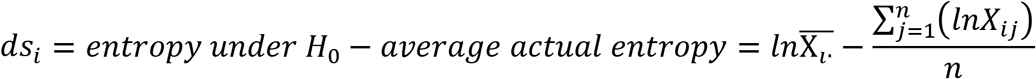

which captures the degree of disorder or randomness of gene expression^37^. Given that genes under *H*_0_ (non-variable genes) account for the major proportion, we fit the relationship between and 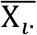 and *average actual entropy*, and calculate corresponding residual as *ds*_*i*_ to imp rove the per formance (Fig. 1b, c). The significance of *ds* is estimated based on a normal distribution approximation and is adjusted using Benjamini-Hochberg method. We also extended such procedure to full-length datasets and found that our approach consistently outperformed other gene selection methods (Fig. 1f,h and Supplementary Fig. 4).

### Data simulation

We simulated droplet datasets with NB distribution. Mean gene abundance levels X were sampled from the log-normal distribution:

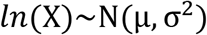

with parameters μ = 0 and σ = 2. The number of transcripts for each gene were drawn from:

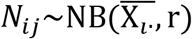

For each simulated dataset, the dispersion parameter r (r = *α*)^41^ was set to a fixed value, ranging from 2 to 10 (Supplementary Fig. 1). In addition, we simulated full-transcript datasets with ZINB distribution. The dropout rates for each gene was modeled with the sigmoid function^42^:

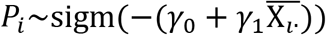

with parameters *γ*_0_ = −1.5 and 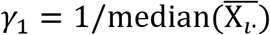. Each simulated scRNA-seq dataset contained 20,000 genes and 2,000 cells (Supplementary Fig. 2).

Differentially expressed genes were added in a fraction of cells (1%-50%, Supplementary Figs. 1 and 2), with fold changes sampled from the log-normal distribution (μ = 0 and σ = 2).

### HVG, Gini and M3Drop

The HVG method^11^ identifies variable genes by comparing the coefficient of variation squared (CV^2^) to a local regression trend, and was implemented with the ‘BrenneckeGetVariableGenes’ function in the M3Drop^14^ package. In the Gini index model proposed in GiniClust^13^, a gene is considered as informative if its Gini is higher than expected from the maximum observed expression. In addition, M3Drop uses dropout rates for variable gene selection and was implemented with the ‘M3DropFeatureSelection’ function in the M3Drop package.

### Cross-validation experiments and gene reproducibility

To illustrate the performance of *S*-*E* model in real datasets (Supplementary Table 1), we performed cross-validation experiments using RF^18^ (with 200 trees) from the python module sklearn^29^, with 70% of the cells randomly selected from the original sample as reference and the remaining 30% as query set. The classification accuracy was measured as the percentage of query cells that were assigned the correct label. We calculated the reproducibility by intersecting the corresponding sets of variable genes as:

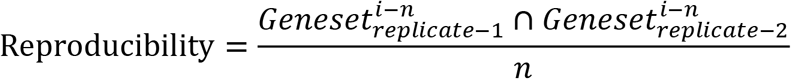

where *i* denotes the gene selection method adapted and *n* is the number of top ranked variable genes.

### ROGUE calculation

By taking advantage of the wide applicability of *S*-*E* model to scRNA-seq data, we introduce the statistic ROGUE to measure the purity of a cell population as:

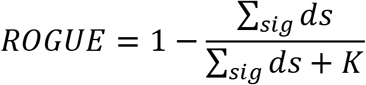

where *K* is an important parameter that constrains the ROGUE value between 0 and 1. A cell population with no significant ds for all genes will receive a ROGUE value of 1, while a population with maximum summarization of significant ds is supposed to yield a purity score of ~0. We reasoned that Tabula Muris^2^ can be considered as such a plausible reference dataset with maximum ∑_*sig*_ *ds* because it comprises cells from 20 organs, and thus represents a highly heterogeneous population. Motivated by the definition of Michaelis constant in Michaelis-Menten equation, the default value of *K* is set to one-half summarized significant ds of Tabula Muris dataset. In this way, ROGUE will receive a value of 0.5 when summarized ds is equivalent to one-half of the maximum, and hence provides a universal metric for assessing the purity of given cell clusters. The default value of *K* is set to 45 and 500 for droplet-based and full-length based datasets, respectively. The *K* value can also be determined in a similar way by specifying a different reference dataset in particular scRNA-seq data analyses.

### Silhouette coefficient

To assess the differences of simulated replicates and the separation of different cell clusters, we calculated the silhouette width^7^, which is the ratio of within-cluster to inter-cluster dissimilarity. Let *a*(*i*) denote the average dissimilarity of cell *i* to all other cells of its cluster A, and let *b*(*i*) denote the average dissimilarity of cell *i* to all data points assigned to the neighboring cluster, whose dissimilarity with cluster A is minimal. The silhouette width for a given cell *i* is defined as:

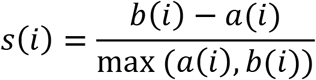

A high *s*(*i*) va lue su ggests that the cell *i* is wel l assigned to its own cluster but poorly assigned to neighboring clusters.

### Sequencing depth simulation

Sequencing depth can vary significantly across cells and thus contributes to a substantial technical confounder in scRNA-seq data analysis. To illustrate that ROGUE is robust to sequencing depth, we generated simulated populations, each consisting of two replicates with only differences in sequencing depth (Fig. 4d and Supplementary Fig. 7a). In each simulation, we varied the sequencing depth of the two replicates as:

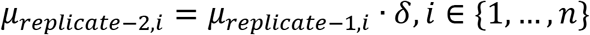

where n is the number of genes, *μ* is the mean expression level and *δ* ∈ {2, 5, 10, 20, 50, 70, 100}.

### Generation of simulated cell types

To demonstrate the potential for ROGUE to guide single cell clustering, we used NB model as aforementioned to simulate different scRNA-seq datasets, each consisting of three cell types A, B and C (1,000 cells × 10,000 genes each), where A and B were similar subtypes. For the three scenarios shown in Fig. 3a and Supplementary Fig. 8a,d, we introduced 500, 1000 and 800 varied genes between cell type A and cell type B/C respectively, with fold changes drawn from the log-normal distribution (μ=0 and σ=2). In addition, we simulated 100, 100 and 120 highly variable genes between cell type B and C respectively, with fold changes sampled from a log-normal distribution with μ=0 and σ=1. The results were visualized using t-distributed Stochastic Neighbor Embedding (t-SNE)^43^.

### Processing and analysis of the fibroblast and B cell datasets

We filtered out low-quality cells with either less than 600 expressed genes, over 25,000 or below 600 UMIs. After filtration, a total of 4,291 B cells and 1,465 fibroblasts were remained. We further normalized the gene expression matrices using regularized negative binomial regression^44^ with the SCTransform function in Seurat^25^. The top 3000 genes with maximal Pearson residual were used for PCA analysis. To remove batch effects between donors, we performed batch correction using BBKNN^45^ with the first 50 PCs. Using the leiden clustering approach implemented in scanpy^46^, each cell cluster was identified by its principle components. This yielded 11 fibroblast subtypes and 7 B cell subtypes as shown in Fig. 3d and Fig. 4b, which were visualized in 2D projection of UMAP^47^ with default parameters.

### Pathway and TCGA data analysis

To characterize and detect the pathway signals in specific fibroblast subtypes, we performed pathway analyses using hallmark pathways from the molecular signature database^48^ with GSVA^49^. The TCGA LUAD and LIHC data were used to investigate the prognostic effect of 13 signature genes (Supplementary Table 5) derived from B_C2_ACTB. To eliminate the effects of different B cell proportions, we normalized the mean abundance level of these 13 marker genes by the expression of MS4A1 gene, and performed subsequent statistical analyses using GEPIA2^50^ with default parameters.

### Data and Software Availability

The datasets used in our study are listed in Supplementary Tables 1-3, along with download links or GEO accession numbers. Our approach is implemented in an open-source R toolkit ROGUE (https://github.com/PaulingLiu/ROGUE) and is freely available to users.

## Acknowledgments

We thank Y. He and D. F. Wang for valuable discussions on this study. This project was supported by Beijing Advanced Innovation Centre for Genomics at Peking University, Key Technologies R&D Program (2016YFC0900100), National Natural Science Foundation of China (31530036 and 91742203).

## Author contributions

Z.Z. conceived this study. C.L. and B.L. designed the *S*-*E* model. B.L. introduced the algorithm of ROGUE, performed the benchmark testing, analyzed the data and developed the R package. Z.L. and X.R. assisted with method development. B.L., C.L. and Z.Z. wrote the manuscript with all the authors’ inputs.

## References

1. Aran, D., Hu, Z. & Butte, A. J. xCell: digitally portraying the tissue cellular heterogeneity landscape. Genome Biol. 18, 220 (2017).

2. Schaum, N. et al. Single-cell transcriptomics of 20 mouse organs creates a Tabula Muris. Nature 562, 367–372 (2018).

3. Tirosh, I. et al. Dissecting the multicellular ecosystem of metastatic melanoma by single-cell RNA-seq. Science 352, 189 (2016).

4. Li, L. et al. Single-Cell RNA-Seq Analysis Maps Development of Human Germline Cells and Gonadal Niche Interactions. Cell Stem Cell 20, 858–873.e4 (2017).

5. Zhang, L. et al. Lineage tracking reveals dynamic relationships of T cells in colorectal cancer. Nature 564, 268–272 (2018).

6. Tian, L. et al. Benchmarking single cell RNA-sequencing analysis pipelines using mixture control experiments. Nat. Methods 16, 479–487 (2019).

7. Rousseeuw, P. J. Silhouettes: A graphical aid to the interpretation and validation of cluster analysis. J. Comput. Appl. Math. 20, 53–65 (1987).

8. Zhang, J. M., Fan, J., Fan, H. C., Rosenfeld, D. & Tse, D. N. An interpretable framework for clustering single-cell RNA-Seq datasets. BMC Bioinformatics 19, 93 (2018).

9. Sade-Feldman, M. et al. Defining T Cell States Associated with Response to Checkpoint Immunotherapy in Melanoma. Cell 175, 998–1013.e20 (2018).

10. Soneson, C. & Robinson, M. D. Bias, robustness and scalability in single-cell differential expression analysis. Nat. Methods 15, 255 (2018).

11. Brennecke, P. et al. Accounting for technical noise in single-cell RNA-seq experiments. Nat. Methods 10, 1093 (2013).

12. Trapnell, C. et al. Differential analysis of gene regulation at transcript resolution with RNA-seq. Nat. Biotechnol. 31, 46 (2012).

13. Jiang, L., Chen, H., Pinello, L. & Yuan, G.-C. GiniClust: detecting rare cell types from single-cell gene expression data with Gini index. Genome Biol. 17, 144 (2016).

14. Andrews, T. S. & Hemberg, M. M3Drop: dropout-based feature selection for scRNASeq. (2018). doi:10.1093/bioinformatics/bty1044

15. Macosko, E. Z. et al. Highly Parallel Genome-wide Expression Profiling of Individual Cells Using Nanoliter Droplets. Cell 161, 1202–1214 (2015).

16. Grün, D., Kester, L. & van Oudenaarden, A. Validation of noise models for single-cell transcriptomics. Nat. Methods 11, 637 (2014).

17. Risso, D., Perraudeau, F., Gribkova, S., Dudoit, S. & Vert, J.-P. A general and flexible method for signal extraction from single-cell RNA-seq data. Nat. Commun. 9, 284 (2018).

18. Breiman, L. Random Forests. Mach. Learn. 45, 5–32 (2001).

19. Papalexi, E. & Satija, R. Single-cell RNA sequencing to explore immune cell heterogeneity. Nat. Rev. Immunol. 18, 35 (2017).

20. Zheng, G. X. Y. et al. Massively parallel digital transcriptional profiling of single cells. Nat. Commun. 8, 14049 (2017).

21. Guo, X. et al. Global characterization of T cells in non-small-cell lung cancer by single-cell sequencing. Nat. Med. 24, 978–985 (2018).

22. Zilionis, R. et al. Single-Cell Transcriptomics of Human and Mouse Lung Cancers Reveals Conserved Myeloid Populations across Individuals and Species. Immunity 50, 1317–1334.e10 (2019).

23. Collin, M. & Bigley, V. Human dendritic cell subsets: an update. Immunology 154, 3–20 (2018).

24. Dutertre, C.-A. et al. Single-Cell Analysis of Human Mononuclear Phagocytes Reveals Subset-Defining Markers and Identifies Circulating Inflammatory Dendritic Cells. Immunity 51, 573–589.e8 (2019).

25. Stuart, T. et al. Comprehensive Integration of Single-Cell Data. Cell 177, 1888–1902.e21 (2019).

26. Lambrechts, D. et al. Phenotype molding of stromal cells in the lung tumor microenvironment. Nat. Med. 24, 1277–1289 (2018).

27. Kalluri, R. The biology and function of fibroblasts in cancer. Nat. Rev. Cancer 16, 582 (2016).

28. Elyada, E. et al. Cross-Species Single-Cell Analysis of Pancreatic Ductal Adenocarcinoma Reveals Antigen-Presenting Cancer-Associated Fibroblasts. Cancer Discov. 9, 1102 (2019).

29. Fabian, P. & Gaël, V. Scikit-learn: Machine learning in Python. J. Mach. Learn. Res. 12, 2825–2830 (2011).

30. Zhang, Q., He, Y., Luo, N. & Zhang, Z. Landscape and dynamics of single immune cells in hepatocellular carcinoma (unpublished).

31. Ohkubo, Y. et al. A Role for c-*fos*/Activator Protein 1 in B Lymphocyte Terminal Differentiation. J. Immunol. 174, 7703 (2005).

32. Kang, H. M. et al. Multiplexed droplet single-cell RNA-sequencing using natural genetic variation. Nat. Biotechnol. 36, 89–94 (2018).

33. Ahmed, R. et al. A Public BCR Present in a Unique Dual-Receptor-Expressing Lymphocyte from Type 1 Diabetes Patients Encodes a Potent T Cell Autoantigen. Cell 177, 1583–1599.e16 (2019).

34. Tedoldi, S. et al. Jaw1/LRMP, a germinal centre-associated marker for the immunohistological study of B-cell lymphomas. J. Pathol. 209, 454–463 (2006).

35. Huang, M. et al. SAVER: gene expression recovery for single-cell RNA sequencing. Nat. Methods 15, 539–542 (2018).

36. Zappia, L., Phipson, B. & Oshlack, A. Splatter: simulation of single-cell RNA sequencing data. Genome Biol. 18, 174 (2017).

37. Li, C. et al. SciBet: a fast classifier for cell type identification using single cell RNA sequencing data. bioRxiv 645358 (2019). doi:10.1101/645358

38. Bacher, R. et al. SCnorm: robust normalization of single-cell RNA-seq data. Nat. Methods 14, 584–586 (2017).

39. Lun, A. T. L., Bach, K. & Marioni, J. C. Pooling across cells to normalize single-cell RNA sequencing data with many zero counts. Genome Biol. 17, 75–75 (2016).

40. Vallejos, C. A., Marioni, J. C. & Richardson, S. BASiCS: Bayesian Analysis of Single-Cell Sequencing Data. PLoS Comput. Biol. 11, e1004333–e1004333 (2015).

41. Lloyd-Smith, J. O. Maximum likelihood estimation of the negative binomial dispersion parameter for highly overdispersed data, with applications to infectious diseases. PloS One 2, e180–e180 (2007).

42. Büttner, M., Miao, Z., Wolf, F. A., Teichmann, S. A. & Theis, F. J. A test metric for assessing single-cell RNA-seq batch correction. Nat. Methods 16, 43–49 (2019).

43. Maaten, L. van der & Hinton, G. E. Visualizing Data using t-SNE. in (2008).

44. Hafemeister, C. & Satija, R. Normalization and variance stabilization of single-cell RNA-seq data using regularized negative binomial regression. bioRxiv 576827 (2019). doi:10.1101/576827

45. Polański, K. et al. BBKNN: fast batch alignment of single cell transcriptomes. Bioinformatics (2019). doi:10.1093/bioinformatics/btz625

46. Wolf, F. A., Angerer, P. & Theis, F. J. SCANPY: large-scale single-cell gene expression data analysis. Genome Biol. 19, 15 (2018).

47. Becht, E. et al. Dimensionality reduction for visualizing single-cell data using UMAP. Nat. Biotechnol. 37, 38 (2018).

48. Subramanian, A. et al. Gene set enrichment analysis: A knowledge-based approach for interpreting genome-wide expression profiles. Proc. Natl. Acad. Sci. 102, 15545 (2005).

49. Hänzelmann, S., Castelo, R. & Guinney, J. GSVA: gene set variation analysis for microarray and RNA-seq data. BMC Bioinformatics 14, 7–7 (2013).

50. Tang, Z., Kang, B., Li, C., Chen, T. & Zhang, Z. GEPIA2: an enhanced web server for large-scale expression profiling and interactive analysis. Nucleic Acids Res. 47, W556–W560 (2019).

